# African Green Monkeys Respond to Synthetic Aβ Oligomers with Persistent Alzheimer’s-like Activation

**DOI:** 10.1101/2025.10.08.678256

**Authors:** Bianca R.P. Brown, Xiaoting Li, Monica R. Grasty, Isabel R. Lopez, Sara Dzigurski, Maya N. Geradi, Michael R. Weed, John D. Elsworth, Matthew Lawrence, Gamze Gürsoy, Andrew D Miranker

**Affiliations:** Department of Molecular Biophysics and Biochemistry Yale University, New Haven, CT 06511, USA; Department of Chemical and Environmental Engineering Yale University, New Haven, CT 06511, USA; Department of Biomedical Informatics, Columbia University, New York, NY 10032, USA; New York Genome Center, New York, NY 10013, USA; Department of Computer Science, Columbia University, New York, NY 10032, USA; Virscio, Inc. New Haven, CT 06511, USA

**Keywords:** Non-human primate, LC-MS, Neurodegeneration, Longitudinal disease model, Liquid biopsy

## Abstract

Wild African green monkeys (AGMs) provide a promising alternative to congenic rodent models because of their closer evolutionary relationship to humans and natural genetic variation. They share key physiological and biochemical traits with humans, including lifespan, neuroanatomy, vascular structure, and inflammatory responses. Unlike rodents, AGMs naturally develop Alzheimer’s-like amyloid-β (Aβ) plaques and tau tangles with age. Immunohistochemical studies further show that AGMs inoculated with synthetic Aβ oligomers (AβO) exhibit hyperphosphorylated tau and neuroinflammation one year later, in the absence of overt neurodegeneration. The AGM body size permits collection of cerebrospinal fluid (CSF) and CSF derived extracellular vesicles (EV) from living individuals, which are key sources of Alzheimer’s disease biomarkers that can be monitored during disease progression. Here, we evaluate AβO treated AGMs at the systems level using proteomics of CSF and phosphatidylserine affinity isolated EVs (EVps). We optimized a workflow to obtain paired CSF and EVps proteomics from <1 mL volumes, i.e. comparable to human liquid biopsy. Our measurements reveal robust, persistent AD-like responses at the biochemical level without overt loss of cognitive function. As such, these findings in AGMs suggest potential alternatives for disease tracking or point to protective mechanisms for limiting disease progression in AD.

**Highlights:**

- Dual proteomics of African green monkeys transiently challenged with synthetic Aβ oligomers (AβO)
- Phosphotidylserine (TIM4) based workflow enables CSF and EV profiling using clinical volumes
- One year post-AβO: vascular-inflammatory pathways rise; neuronal-axonal pathways fall
- AβOs drive human-AD-like proteome shifts on time scales shorter than cognitive decline

**In brief:** Wild African green monkeys (AGMs) offer a translational model for early Alzheimer’s biology, with physiology more similar to humans. We transiently exposed AGMs to synthetic Aβ oligomers and, 12 months later, profiled paired proteomes from whole CSF and a CSF subcompartment enriched for extracellular vesicles. Despite no overt cognitive decline, AGM proteomes showed persistent Alzheimer’s-like remodeling, particularly in vascular, inflammatory, and neuronal systems. Parallel analysis of the CSF subcompartment revealed proteins and pathways under-represented in bulk CSF, sharpening disease-relevant signals and candidate biomarkers. This systems-level, longitudinal study establishes AGMs as a powerful platform for liquid biopsy discovery and illuminates basic biology of molecular responses to soluble Aβ oligomers accompany and potentially protect against neurodegeneration.

## INTRODUCTION

The continued failure of Alzheimer’s disease (AD) drug development is largely driven by the lack of animal model systems that faithfully mirror the key biological and physiological features of the human disease. AD is clinically defined by progressive memory loss and cognitive decline, and pathologically by amyloid-β (Aβ) plaques, tau tangles, neuroinflammation, and neuronal loss. Trials targeting Aβ, such as gamma-secretase inhibitors, have consistently failed with suggestions that Aβ initiates pathology but tau drives progression^1,2^. Evidence now points to soluble Aβ oligomers (AβO), rather than plaques, as the toxic species disrupting synapses and memory^3^. The implication of AβO has redirected the field’s focus toward the full trajectory of AD, spanning the early formation of AβO^3^ through downstream events such as tau tangle accumulation, and neuroinflammation^4^. Recognizing this complexity has created a demand for models that can encompass both the initiating events driven by Aβ oligomers and the downstream cascades leading to neurodegeneration in humans.

The African green monkey (AGM) offers a compelling alternative to traditional rodent models for AD research. Non-human primates have been used for translation because they are evolutionarily closer and more physiologically similar to humans than other model systems^5–7^. AGMs diverged from the human lineage approximately 25–30 million years ago, compared to ∼90–100 million years for mice^8^. Protein analyses of brain tissue show that aged AGMs naturally exhibit Alzheimer’s-like pathology, with both Aβ40 and Aβ42 levels increasing with age^9^. Rodent systems capture this trajectory poorly, as many of these downstream processes are absent or only weakly represented^10^. Rodent models respond neurologically to Aβ, validating its role as an initiator of pathology, but they fail to capture downstream effects^11^. Tau, a key mediator of neurodegeneration, does not undergo pathological changes in rodents.

AGM also has the advantage of being a prodromal model, providing early insights into AD prior cognitive decline. Although aged AGMs develop Aβ plaque deposition and increased phosphorylated tau, they do not progress to full Alzheimer’s pathology, lacking widespread tau tangles and neurodegeneration^6,12^. Recent studies show that treatment with AβO administration induces downstream tau hyperphosphorylation in the medial temporal lobe, a hallmark of early-stage AD out to at least 3 months post AβO exposure^13^. Despite showing this neuroinflammation and tau pathology, AGMs exhibited no overt neurological decline over 12 months (Weed unpublished data) making them a promising model for studying early AD pathogenesis. However, because brain tissue can only be collected post-mortem, disease progression cannot be tracked directly in living AGMs. Biofluids such as cerebrospinal fluid (CSF) provide an alternative window into central nervous system processes, enabling repeated sampling and longitudinal monitoring of AD-related changes from the earliest stages.

CSF and its associated extracellular vesicles (EVs) capture different but complementary biological signatures of Alzheimer’s disease, providing synergistic insights into disease progression and biomarker development. Clinical symptoms of Alzheimer’s typically begin in the hippocampus before spreading to other brain regions, leading to progressive functional decline. This spread reflects the propagation of a disease through cell-cell communication^14^. EVs may contribute to this communication by facilitating the transfer of pathogenic proteins across brain regions. At the same time, EVs serve as quantifiable carriers of early and region-specific molecular signals in CSF. As CSF circulates through the central nervous system, it exchanges proteins with brain tissue, and many of these proteins are also packaged into EVs. By enriching a subset of proteins, EVs have the potential to reveal low-abundance biomarkers that remain undetectable in bulk CSF analyses^15^. Together, CSF and EVs provide a more complete view of the communication landscape in Alzheimer’s disease, capturing both shared and unique signals that can inform biomarker discovery.

Here, we evaluate the feasibility of using the AGM model to capture Alzheimer’s-related changes at the proteome scale in a minimally invasive manner. In this pilot study for biomarker discovery, we assess whether AβO treatment induces proteome shifts in whole CSF and CSF-derived EVs that may be informative for future human and animal model based studies (Figure 1). While several studies have used proteomics of CSF to identify AD-associated pathways^16–20^, there have been increased interest in CSF derived EVs ^21–23^. As such, our goals were to: (1) profile the proteomes of whole CSF and CSF-derived EVs using methods we optimized for small volumes; (2) compare proteomic changes between AβO-treated and control AGMs; and (3) test whether AβO-responsive proteins in AGMs align with human Alzheimer’s disease signatures.

**Figure 1.**
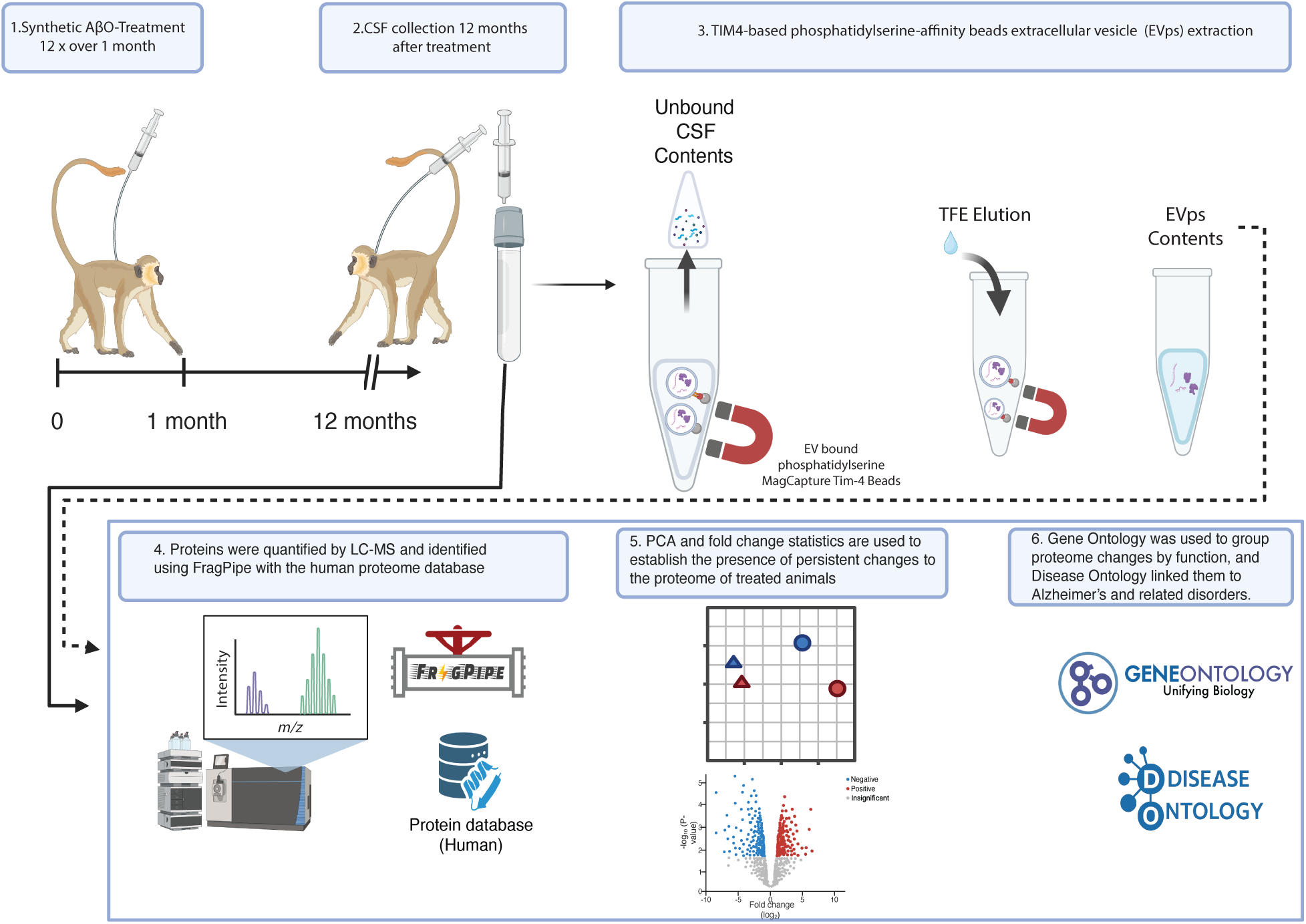
Study design and analysis workflow. (1) Monkeys were administered AβO three times per week for one month, and (2) twelve months after the final dose cerebrospinal fluid (CSF) was collected from the cisterna magna of both AβO-treated and vehicle-control individuals. (3) EVps were extracted from CSF using phosphoserine MagCapture beads, and (4) raw mass spectrometry scans were processed to identify proteins and peptides. (4) Data analysis included differential abundance testing at the protein level, followed by (5) gene ontology (GO) enrichment and disease ontology (DO) association to interpret functional and disease-related signatures.

## RESULTS

### Study design

We challenged African green monkeys with synthetic Aβ oligomers (AβO) or vehicle (Ham’s F12 and saline) following established protocol (Wakeman et al., 2022). Twelve months after the final dose, CSF was collected for paired profiling of whole CSF and the phosphatidylserine-positive nanoparticle fraction (EVps) using commercial TIM4-based affinity beads we optimized for use with small volumes (Figure 1). The use of vehicle control and year wait post AβO challenge was designed to isolate AβO-specific effects and diminish potential for contributions from repeated intrathecal delivery. Proteomic analysis, presented below, revealed preclinical, Alzheimer’s-like shifts, with EVps partitioning highlighting biology not evident in bulk CSF, underscoring the model’s value for biomarker discovery.

In total, we were able to acquire data from 4 whole CSF samples (2 AβO, 2 vehicle) and 6 EVps samples (3 AβO, 3 vehicle). Tryptic digest followed by liquid chromatography mass spectrometry (LC–MS) to identify and perform label-free quantification of the proteome of each sample. We used FragPipe/MSFragger to identify peptides^24^ and mapped them to the human proteome, both to determine whether AGMs express proteins that serve as human relevant markers and to provide a comprehensive reference for downstream ontology analyses. In order to enable comparative analysis, we normalized all MS intensities collected by IonQuant DDA (see methods)^24^. We then assessed protein intensity differences between AβO and vehicle samples of CSF and EVps using a workflow that combined log₂-normalized PCA with mean-quantifier analysis (see Methods). We mapped the resulting differentially abundant proteins to Gene Ontology and Disease Ontology categories to reveal system-level shifts in the proteome.

### CSF and EVps proteomes emphasize functionally distinct compartments with overlapping proteins

CSF and EVps share proteins but also differ, with EVps representing a selective subset that enriches for signals missed in whole CSF. We determined which proteins were unique and which were shared by assessing their presence or absence in each compartment, defining presence as proteins with intensities above 0. Across all samples, we identified 738 unique proteins. Of these, 214 were shared between the whole CSF and its EVps fraction, representing ∼32% of the total CSF proteome and 78% of the EVps proteome (Figure 2A & S1A; Table S1). This suggests EVps may enrich for proteins that whole CSF analyses may overlook.

**Figure 2.**
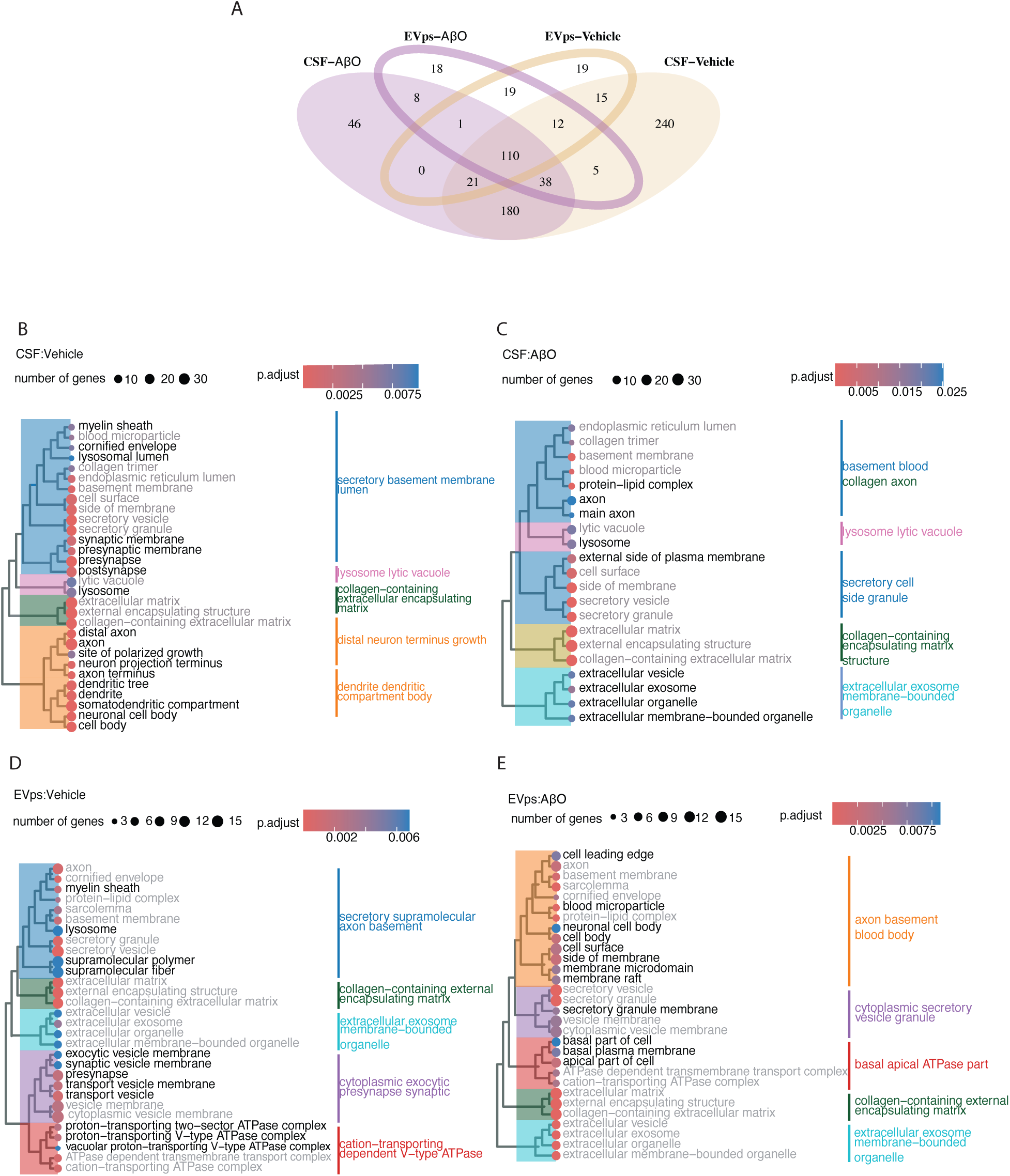
Overview of proteins across whole CSF and EVps. (A) Overlap of proteins identified in CSF and EVps samples from both AβO-treated and vehicle-control individuals. Top 30 enriched Gene Ontology (Cellular Component) terms, ranked by adjusted p-values and clustered by similarity, are shown for (B) CSF-AβO, (C) CSF-control, (D) EVps-AβO, and (E) EVps-vehicle samples. Grayed functions are shared within subcompartments. See Figure S5 for Gene Ontology terms associated with biological processes and molecular functions.

Although the EVps compartment represented a subset of whole CSF, it was clearly distinct, revealing proteins not detected in CSF and showing divergent intensity patterns for proteins shared between the two compartments. First, 54 of 738 proteins were observed only in the EVps compartment. These proteins were likely detectable because of their selective enrichment and the elimination of CSF proteins that might otherwise have obscured their presence when measured by LC-MS.

### Overlapping proteins show distinct abundance profiles in CSF and EVps

In CSF, shared proteins were disproportionately drawn from the highest abundance fraction: their mean log₁₀ intensity (8.7 ± 0.8) was higher than the average for all proteins (8.1 ± 0.8) and the CSF-unique subset (7.8 ± 0.6; Figure S1B). In EVps, however, overlapping proteins showed no such enrichment. Their mean intensities (7.8 ± 0.7) were nearly identical to those of the total EVps proteome (7.3 ± 0.7) and the EVps unique subset (7.1 ± 0.6; Figure S1C). This indicates that EVps do not simply carry over the most abundant CSF proteins but instead represent a distinct, selectively packaged subset.

Gene ontology results confirmed that we captured information representative of the central nervous systems (CNS) of the AGMs. EVps and CSF shared a common set of functions, including extracellular exosome and extracellular vesicle pathways, membrane-bounded organelles, secretory vesicles and granules, collagen containing extracellular matrix and basement membrane terms, and axon associated compartments such as the axon and axon basement (Figure 2B-D; Figure S1C). Beyond this overlap, CSF uniquely contained degradative and structural remodeling functions, including lysosome and lytic vacuole, neuronal soma and dendrite, distal neuron terminus, neuronal projection, postsynapse, growth cone, and polarized growth sites (Figure 2B,C). By contrast, EVps uniquely contained synaptic and transport-focused functions, including presynapse and synaptic vesicle membrane, supramolecular fibers such as the myelin sheath and protein–lipid complexes, multiple ATPase complexes (proton-transporting ATPase, vacuolar V-type ATPase, and ATPase-dependent transmembrane transport), and polarity-linked terms such as cell leading edge, basal plasma membrane, and membrane rafts or microdomains (Figure 2D,E). These differences indicate that while both compartments share a baseline of vesicle secretion, extracellular matrix, and axonal functions, EVps were distinguished by synaptic trafficking and proton-pumping machinery, whereas CSF is enriched for degradative pathways and large-scale neuronal remodeling.

Both PCA and differential abundance analyses demonstrated that CSF and EVps proteomes were distinct, highlighting EVps were not simply scaled-down versions of CSF. To assess global proteome differences between AβO-treated and vehicle controls, we performed principal component analysis (PCA) on the normalized log_10_ intensities detected in both compartments. This analysis revealed clear separation of CSF and EVps along PC1, which explained 56% of the total variance (Figure 3A).

**Figure 3.**
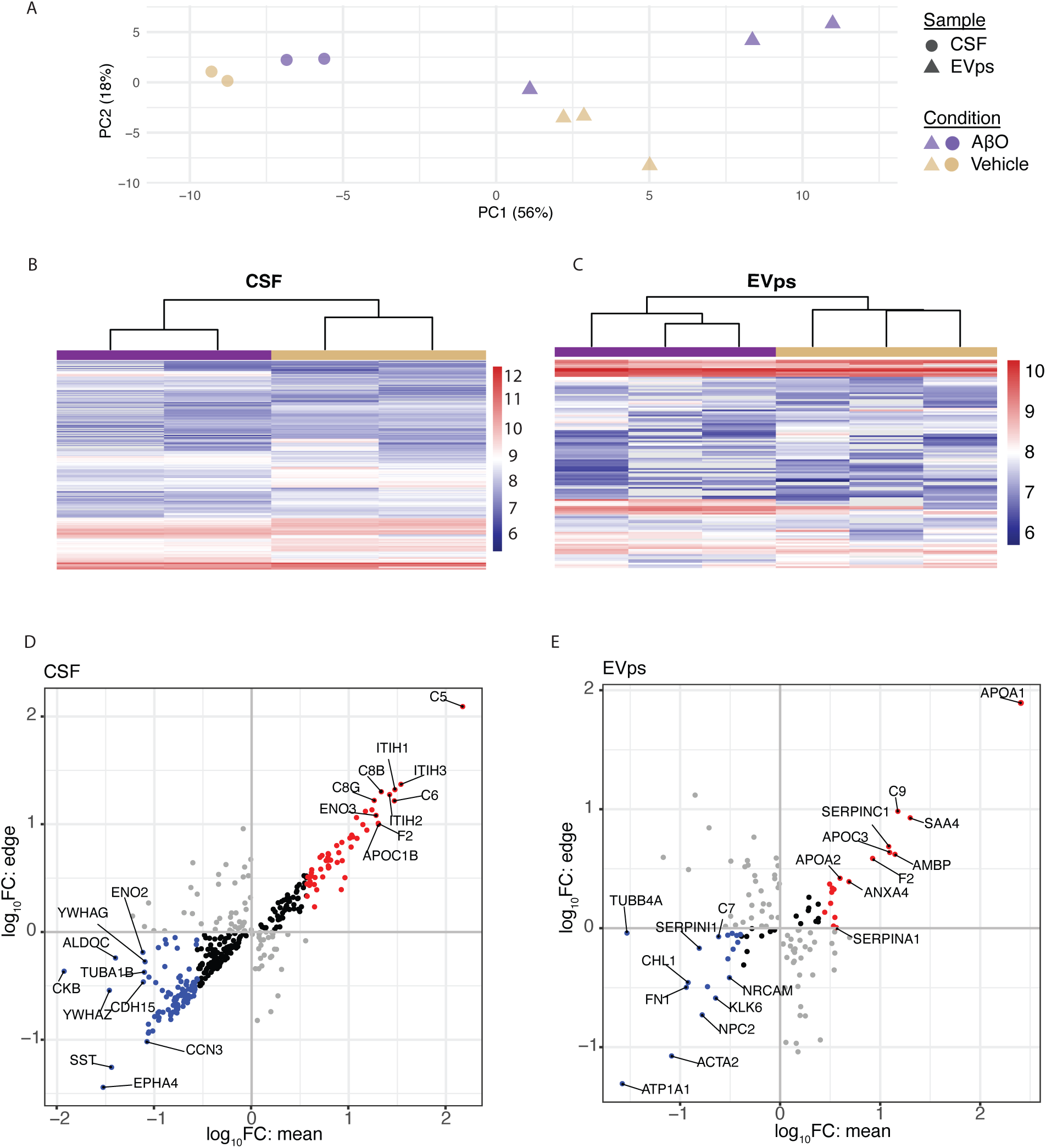
Global proteome differences between AβO- and vehicle-treated individuals. (A) Principal component analysis (PCA) of log₁₀-transformed protein intensities for whole CSF and EVps samples. (B-C) Heatmaps of log₁₀-transformed protein intensities for whole CSF (B) and EVps (C) samples, with hierarchical clustering of individual samples shown at the top and treatment conditions (AβO or vehicle) indicated. Only proteins detected in at least 3 of 4 CSF samples or 4 of 6 EVps samples were included to minimize missing data and ensure robust comparisons. (D–E) Differential abundance analyses for whole CSF (D) and EVps (E), showing log₁₀ fold change based on group means (x-axis; log₁₀FC_mean) versus fold change between the closest treatment and control values (y-axis; log₁₀FC_edge). Black points represent “regulated core,” where all treatment values are consistently higher or lower than controls (i.e., log₁₀FC_mean and log₁₀FC_edge share the same sign). The top 10 up- and down-regulated proteins within regulated core groups are labeled.

Overall, these results demonstrate that our optimized method successfully captured CNS proteins. Moreover, the EVps compartment, while a subset of whole CSF, is clearly distinct. EVps reveals proteins not otherwise observed and, for those shared with CSF, exhibits a different intensity distribution.

### AβO induces compartment- and treatment-specific proteomic remodeling

AβO-treated and vehicle-control samples differed in both the number and composition of proteins in CSF and EVps compartments, indicating that these analyses can capture protein level changes induced by synthetic AβO.

We detected a common set of 110 proteins in all four groups (Figure 2A), suggesting a core proteome was maintained regardless of treatment or compartment. Despite this shared core, each group also exhibited unique proteins, reflecting treatment- and compartment-specific composition (Figure 2A). CSF from vehicle-control animals contained the greatest number of unique proteins (n = 240), followed by CSF from AβO-treated animals (n = 46), EVps from vehicle-controls (n = 19), and EVps from AβO-treated animals (n = 18). Overlap patterns also differed by treatment. For example, 180 proteins were shared between CSF-AβO and CSF-vehicle, while only 38 were shared between EVps-vehicle and CSF-vehicle. These treatment-specific overlaps indicate that AβO not only alters protein composition within each compartment but also shifts the degree of correspondence between CSF and EVps.

PCA and fold change analysis confirmed persistent proteomic differences in treated animals. Within each compartment, AβO-treated and vehicle samples clustered separately, whether in CSF or EVps (Figure 3). Separation was particularly strong in CSF, where intergroup differences exceeded intragroup variation. EVps showed greater within-group variability, and although AβO and vehicle samples generally separated, some overlap reflected subtle sample level heterogeneity (Figure 3A). Separate PCAs for CSF and EVps are shown in Figure S3. Hierarchical clustering of protein intensities further supported treatment-based segregation in both compartments (Figures 3B,C).

To identify proteins most affected by treatment, we calculated fold changes in protein intensities between AβO-treated and vehicle-control groups using a mean-quantifier analysis. First, we compared average protein levels between groups to obtain a mean log₁₀ fold change (log₁₀FC_mean). Second, we calculated an “edge effect” measure (log₁₀FC_edge) by comparing the most extreme values, defined as the highest intensity in one group and the lowest intensity of the same protein in the other group (see Methods). Proteins were considered “regulated core” across AβO-treated and vehicle-control groups if both the mean and edge effect measures pointed in the same direction in both groups. Uniformly regulated core proteins with positive values, enriched in AβO-treated compared to vehicle controls, were classified as upregulated, whereas those with negative values, reduced in AβO-treated samples, were classified as downregulated. Within this regulated core group, we identified 266 proteins in the whole CSF and 50 in EVps samples (Table S3). The top ten upregulated and downregulated proteins in each compartment were ranked by log₁₀FC_mean (Figure 3D,E). Several proteins showed differential abundance in both CSF and EVps. APOA1, APOA2, C9, and SERPINA1 were upregulated in both datasets, while TUBB4A was downregulated in both. These proteins are implicated in Alzheimer’s disease through their roles in lipid metabolism, amyloid clearance, neuroinflammation, and protease regulation.

Beyond these overlaps, the two fractions showed distinct profiles. In CSF, additional upregulated proteins included C3, C5, C8, ENO2, and APOC1B, while downregulated proteins included SST, EPHA4, and CKB. In EVps, unique upregulated proteins included SAA4, ANXA4, and AMBP, whereas downregulated proteins included NPC2, NRCAM, KLK6, and ATP1A1. This highlights a core set of overlapping proteins with consistent directionality, alongside sample-type specific changes.

AβO treatment elicited a two-pronged phenotype. Vascular, inflammatory, and metabolic pathways were increased in ABO-treated individuals were upregulated (Figure 4A,C). While synaptic, axonal, and morphogenetic pathways were downregulated (Figure 4B,D). To dissect these shifts, we performed Gene Ontology (GO) enrichment analysis for Biological Processes (BP) on proteins with higher abundance in AβO-treated versus vehicle-control samples (upregulated) and those with lower abundance (downregulated).

**Figure 4.**
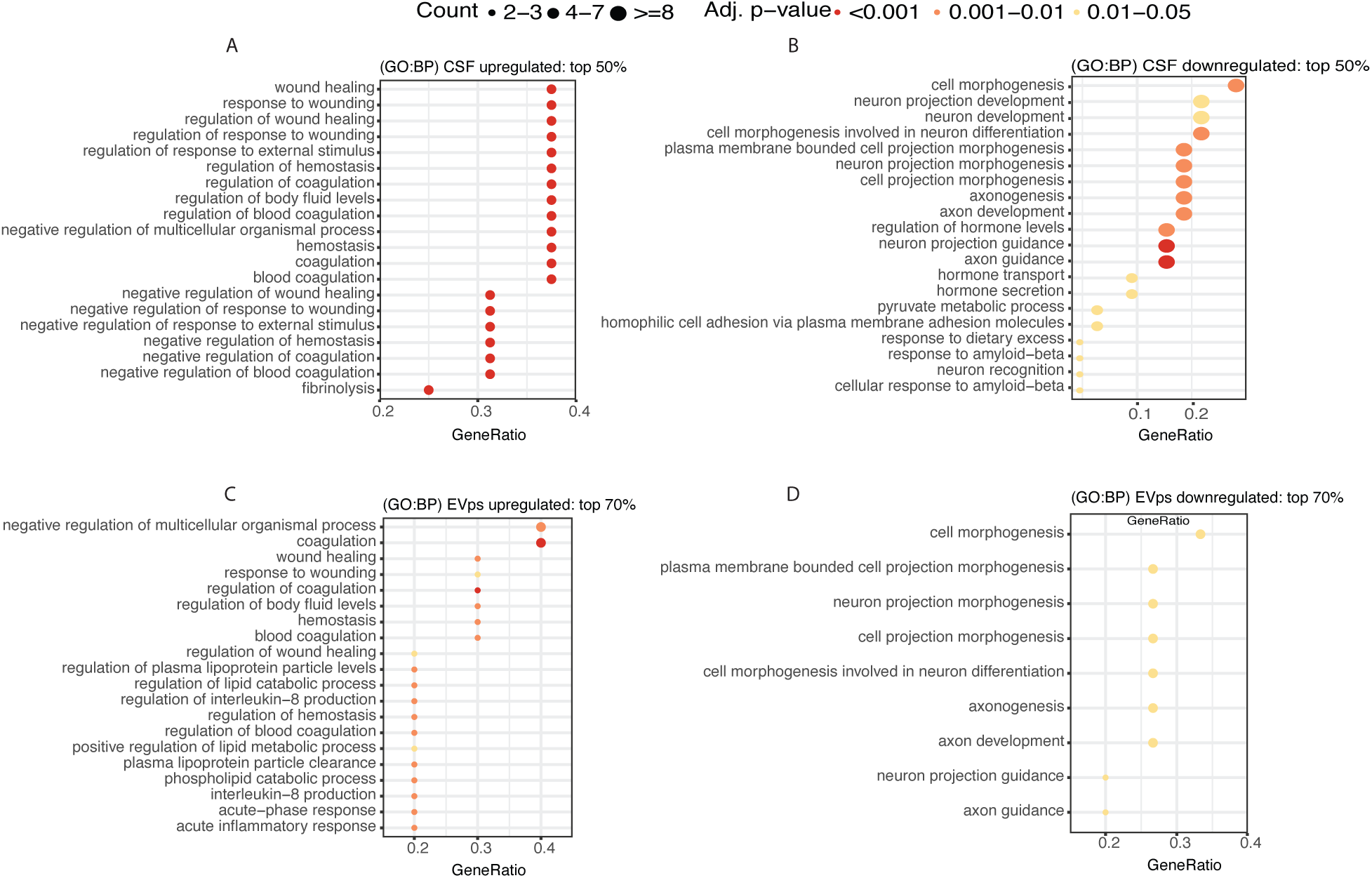
Gene Ontology (GO) enrichment of differentially abundant proteins. Dot plots show the top 20 enriched GO terms ranked by adjusted p-values. For CSF samples, enrichment was computed separately for the top 50% of upregulated proteins (A) and the top 50% of downregulated proteins (B). For EVps samples, enrichment was computed for the top 70% of upregulated (C) and downregulated (D) proteins, reflecting the smaller number of proteins detected.

In CSF, the downregulated proteins in AβO-treated animals were enriched for neuronal pathways, including axon guidance, neuronal differentiation, and cell morphogenesis (Figure 4A; Table S4), indicating a loss of synaptic and axonal functions. In contrast, upregulated proteins in CSF were enriched for vascular processes, including regulation of coagulation, hemostasis, fibrinolysis, wound healing, and response to wounding (Figure 4B). These increases align with Alzheimer’s-related vascular dysfunction, where amyloid–fibrin interactions alter clot structure, impair fibrinolysis, and disrupt cerebrovascular homeostasis.

In EVps, we found that downregulated proteins were enriched for processes related to cell morphogenesis, axon development, and plasma membrane organization (Figure 4D; Table S4), consistent with a loss of neuronal structural and axonal functions. Conversely, upregulated proteins in EVps were enriched for coagulation and wound-healing pathways, as well as acute-phase response, interleukin-8 production, and lipid catabolic processes (Figure 4C). These shifts highlight neuroinflammatory and metabolic changes that complement the vascular signals observed in CSF. These responses closely mirror Alzheimer’s disease hallmarks, including vascular injury, chronic neuroinflammation, impaired lipid metabolism, and progressive loss of neuronal connectivity.

### AβO shifts CSF and EV proteomes toward Alzheimer’s disease hallmarks

Proteins altered by AβO treatment map strongly to Alzheimer’s disease. We investigated Alzheimer’s-specific proteins within the uniformly regulated group using agnostic Disease Ontology (DO) annotations of human genes (Figure 5). AβO treatment shifts the CSF proteome toward Alzheimer’s disease hallmarks by simultaneously increasing inflammatory pathways while reducing neuronal and protective functions. In whole CSF, we identified 26 proteins linked to Alzheimer’s disease, of which 3 were upregulated and 23 were downregulated in AβO-treated samples compared to vehicle-control (Figure 5A). The upregulated proteins reflected a relative gain in inflammatory and lipid-associated responses, including APOA1 (lipid transport and Aβ clearance), C1QB (synaptic pruning and neuroinflammation), and C3 (a complement component linked to Alzheimer’s pathology; Figure 5A; Figure S4 A-C). In contrast, the downregulated proteins represented a relative loss of synaptic, protective, and survival functions, including EPHA4 (synaptic plasticity), TIMP2 (neuroprotection and matrix stability), and PEBP1 (neuronal survival and stress signaling; Figure 5A; Figure S4 D-F).

**Figure 5.**
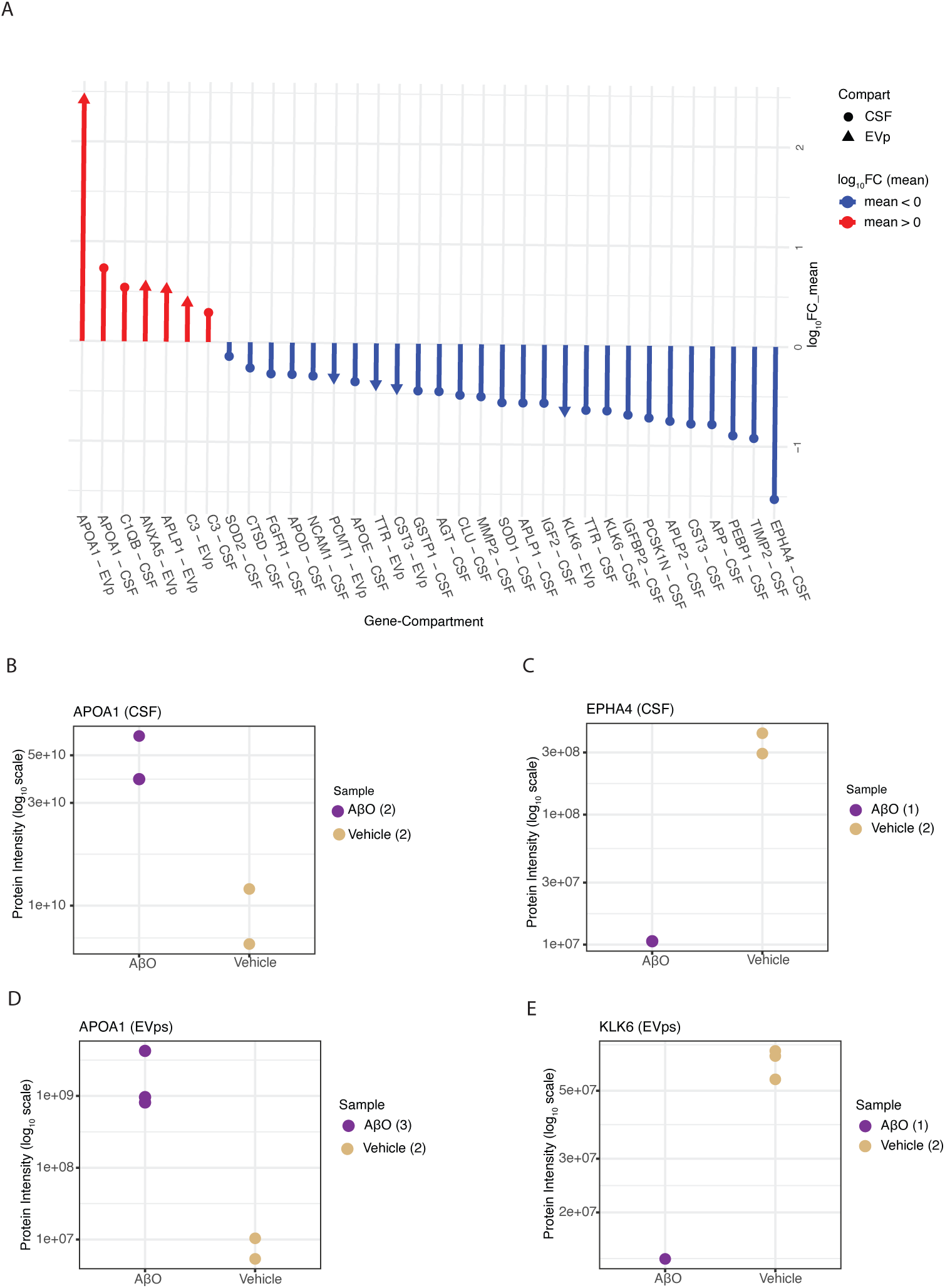
Alzheimer’s Disease-Associated Differentially Abundant Proteins. (A) Line plots showing log₁₀FC_mean values of proteins within the regulated core group that are annotated to Alzheimer’s disease in the Disease Ontology database, for whole CSF (circles) and EVps (triangles). (B, D) Protein intensities of the most (B,D) upregulated and most (C,E) downregulated Alzheimer’s-associated proteins across individual samples, shown separately for whole CSF (B-C) and EVps (D-E).

The altered EV cargo composition in AβO-treated samples suggests that EVps may reflect both increased inflammatory and lipid-associated responses and reduced protective or homeostatic functions, consistent with cellular stress responses associated with amyloid accumulation. In EVps, we identified 8 Alzheimer’s-associated proteins, with 4 upregulated and 4 downregulated in AβO-treated samples compared to vehicle (Figure 5A; Figure S4G-I). The upregulated proteins overlapped with those in whole CSF (APOA1, APLP1, and C3) and included one unique to EVps, ANXA5. The downregulated proteins included PCTM1, TTR, CST3, and KLK6 (Figure 5A; Figure S4J-L). These proteins span diverse functions, including membrane repair (ANXA5), immune signaling (C3), lipid and protein homeostasis (APOA1), and synaptic remodeling (APLP1), highlighting their functional relevance to neurodegenerative processes.

Notably, APOA1 was upregulated in both compartments, but the magnitude was substantially greater in EVps. In CSF, APOA1 increased by ∼8-fold (Figure 5B), whereas in EVps it increased by ∼300-fold (Figure 5D), representing a ∼40-fold stronger effect in EVps. The differences in magnitude of APOA1 highlight the value of including EVps in Alzheimer’s studies. While CSF and EVps show similar directions of regulation for shared proteins, EVps exhibited larger fold-changes, amplifying proteomic differences between conditions and providing a more sensitive readout of disease-associated changes than whole CSF alone.

In addition to Alzheimer’s-specific proteins, differentially abundant proteins were linked to a broader set of disease categories (Figure S5-S6; Table S5). In CSF, upregulated proteins were enriched for immune- and vascular-related diseases, including complement deficiency, blood coagulation disorders, coronary artery disease, and primary immunodeficiency (Figure S5 A-D), while downregulated proteins were enriched for neurodegenerative and protein-misfolding diseases such as tauopathy, Parkinson’s disease, prion disease, synucleinopathy, and macular degeneration (Figure S5 E-H). In EVps, upregulated proteins were associated with metabolic diseases, including lipid metabolism defects, familial hyperlipidemia, and proteinuria (Figure S6 A). Downregulated proteins in EVps were enriched only for amyloidosis (Figure S6 B), indicating a more restricted disease signal than in CSF. Together, these patterns show that AβO treatment produces distinct disease associations across compartments: in CSF, gains are dominated by immune and vascular diseases while losses are linked to neurodegenerative diseases, whereas in EVps, gains are dominated by metabolic diseases and losses are confined to amyloidosis. These associations suggest that AβO-induced proteomic changes extend beyond classical neurodegenerative categories and intersect with immune, vascular, and metabolic disease processes. This network of disease links highlights the systemic nature of Alzheimer’s pathology, reflecting a breakdown of multiple physiological systems rather than an isolated brain disorder.

## Discussion

Identifying animal models that reliably recapitulate Alzheimer’s disease remains a major challenge and a barrier to understanding and ultimately curing the disease. Here, we performed a systems-level assessment of the African green monkey’s display of Alzheimer’s-like physiological changes twelve months after cessation of intrathecal inoculations with synthetic amyloid-beta oligomers. Using only 650 µL of CSF, we performed proteomics on both whole CSF and a subcompartment enriched in phosphotidylserine containing vesicles and other nanoparticles. Together, the whole CSF and EVps subcompartment captured features of prodromal AD pathology. CSF proteome shifts were associated with neuronal injury, while EVps changes highlighted vesicle enriched transport functions in response to treatment. These findings demonstrate the feasibility of dual CSF and EV proteomics from clinically relevant sample volumes and position the AGM as a powerful model for identifying early protein signatures of AD that may be missed in human studies.

AGMs are uniquely suited for longitudinal, biofluid-based studies of AD disease progression, enabling combined evaluation of proteomic signatures in CSF and EVps. CSF is considered superior to post-mortem brain tissue for biomarker discovery in Alzheimer’s disease because it allows repeated sampling across disease stages, whereas brain tissue provides only a single snapshot, often at late-stage pathology, and is subject to variability introduced by post-mortem interval and storage conditions^25^. However, obtaining sufficient CSF volume is a major challenge in rodent models, where total CSF volume is <200 μL (and as low as 40 μL in mice), and long-term sampling is limited by the technical difficulties of catheterization^26,27^. In contrast, nonhuman primates can tolerate intrathecal (ITH) catheters that permit repeated collection of ∼1 mL of CSF, as well as cisternal puncture, providing minimally invasive access to sufficient sample volume^13,28^.

AGMs capture proteomic signatures of AD that rodent models rarely exhibit, even in specialized AβO focused systems. For example, the APP E693Δ-Tg (“Osaka,” APPE693Δ) mouse produces abundant AβO and develops synaptic and cognitive impairments from about 8 months of age, yet it does not form plaques or show tau pathology^29^. In wild-type rats, intracerebroventricular injection of human AβO induces social memory impairment within 21 days^30^. These approaches robustly reproduce selected aspects of AD pathology, but the response is both incomplete and unnaturally rapid. As a result, rodent models provide valuable mechanistic insights but fail to capture the slower, transitional states that define the prodromal stage of Alzheimer’s disease. By contrast, our AGM study reveals a state in which neuropathology develops without corresponding cognitive decline under experimental conditions. These neuropathological changes also arise naturally during AGM aging positioning this model between short-lived rodent systems and chronic human disease^6,12^. In this intermediate space, AGMs offer unique leverage for identifying early biomarkers and testing interventions for AD.

CSF and EVps capture complementary but distinct proteomes, a reflection of their potential to capture different aspects of AD pathology ^15,21,31^. In our hands, in which we performed analysis on matched samples, the two compartments captured both neuronal and vesicular biology albeit with different emphasis. CSF was enriched for neuronal morphology and injury-related signals, highlighting its capacity to reflect neuronal damage and associated immune processes. In synergy, EVps were enriched for protein–lipid complexes, exosome-specific categories, and protein trafficking terms, consistent with their role in reporting ongoing cellular responses within a stressed environment. These functions are similar to that of CSF and EVps in humans^21^. Together, these complementary signatures suggest that dual CSF and EV proteomics can provide a systems-level view of AD pathology, with the ability to link neuronal injury with the cellular responses that accompany it.

The AβO-treated induced shift in the proteome suggests a prodromal Alzheimer’s state, in which cellular stress precedes frank neurodegeneration. Many studies have shown that immune and vascular systems, extracellular and cellular morphology (including neuronal) and regulation of Aβ formation functions are often early signs of the disease^16,32–35^. Broadly, the AβO treated AGMs captured changes in proteins associated with these reported early physiological signs. This underpins our suggestion that our AGM model’s functional proteome response is most closely similar to early stages of AD.

Alongside AGM changes that recapitulate human AD signatures, we also observe alterations in proteins that are infrequently reported in human studies^16–20^. As the model is phylogenetically similar to us, these discrepancies can engaged more usefully and with greater confidence than had they been observed in rodents. First, differences may be a consequence of the AGM represent a longitudinal preclinical model, whereas human studies are never longitudinal and focus on individuals who have progressed to a state of mild cognitive impairment or beyond. In our monkeys, cognitive decline is not apparent. An attractive possibility is that the protein changes we uniquely detect in the AGM model may represent biomarkers with utility for preclinical stages of disease^19,36^. An equally attractive alternative is that the AGM does not display cognitive decline as it has an altered protein expression profile that is neuroprotective, suggesting a roadmap for novel therapeutic targets. Second, the mode of disease induction in the AGM includes the use of young animals and a high degree of insult in the form of injected AβO oligomers. In contrast, Alzheimer’s disease in humans develops over decades and involves complex interactions with aging and comorbid physiological changes. Third, interspecies differences in baseline CSF proteomes (AGM vs. human) may predispose AGMs to give some distinct responses to AβO. I.e. although AGMs are far closer to humans than rodents, they are nevertheless a different family. While we acknowledge these differences, the vast majority of the protein shifts we observed align with human Alzheimer’s-relevant functional changes, making this model especially valuable for probing the transitional period where early stress responses and potential neuronal resilience coexist.

Lipoproteins are a long recognized factor in AD studies and include members such as ApoE for which single nucleotide polymorphisms are strongly associated with disease. Susceptibility. Inclusion of lipoproteins as part of the EVps fraction was therefore paramount to our study^37,38^. Using our small sample optimized phosphatidylserine (PS) affinity bead workflow, we were able to enrich the lipoprotein contribution to our EVps isolates ^39^. Our results revealed lipid metabolism pathways were strongly represented in the PS-captured subcompartment. Lipoproteins such as APOA1, APOA2, and APOC3 were upregulated in AβO-treated EVps, whereas only APOA1 was detected in AβO-treated CSF. Even for APOA1, the magnitude of treatment-associated change was greater in EVps than in CSF. Whether this enrichment reflects the EVs themselves or is enhanced by the PS-based capture remains unclear. Nevertheless, it is clear our approach provides an advantage by revealing lipid-particle-associated features of Alzheimer’s-like pathology that are obscured when measured only using whole CSF.

AβO treatment enriched several neurodegeneration-associated pathways beyond those specific to Alzheimer’s disease. Pathway and disease enrichment analyses here revealed that proteins in CSF fractions were linked to neurological conditions including macular degeneration, prion disease, and synucleinopathies. These results are consistent with previous work suggesting that Alzheimer’s disease emerges from a convergence of multiple pathological processes rather than a single etiology^36^. Notably, among the top enriched disease terms in the EV fraction, only amyloidosis was represented-suggesting a more selective or focused signal. This implies that while specific differentially abundant proteins may vary between monkey and human CSF and EVps, the broader proteome patterns reflect disease relevant biology.

## Conclusion

Our results demonstrate that transient introduction of AβO to the CSF of AGMs induces persistent, AD related shifts in the proteomes of both CSF and EVps compartments. This study establishes a practical framework for small volume, untargeted proteomes to detect early disease signatures in a translational nonhuman primate model. To our knowledge, this is the first study to integrate CSF and CSF-derived EVps proteomes from individual non-human primates, expanding the landscape of Alzheimer’s biomarker discovery beyond brain tissue and blood focused approaches. Importantly, this work positions the African green monkey as a tractable and physiologically relevant system for studying Alzheimer’s like pathology and potentially other neurodegenerative disorders, with the capacity for repeated, longitudinal biofluid sampling. By integrating CSF and EVps, our approach provides a scalable framework for small-volume, multi-compartment proteomics, enabling biomarker discovery not only for Alzheimer’s but across neurodegenerative conditions.

## METHOD DETAILS

### Study system

Our study focused on 6 female African green monkeys (*Chlorocebus aethiops sabaeus*). These monkeys were sourced from a natural free range population historically introduced to St. Kitts, West Indies. In vivo experiments were performed in Virscio’s facilities at the St. Kitts Biomedical Research Foundation, Lower Bourryeau Estate, St. Kitts and Nevis. Studies were approved by the Virscio IACUC and conducted following the National Research Council and AAALAC standards. At the time of initial dosing animals were of normal health and between 6 and 9 years with no detected Alzheimer’s-associated phenotypic traits.

### Experimental design

Adult female monkeys were inoculated with AβO using lumbar intrathecal ports following protocols for surgery and dosing as previously described in Wakeman et al. 2022. Briefly, AGMs were sedated with ketamine (8 mg/kg, IM), given atropine (0.02–0.04 mg/kg, IM), and maintained under isoflurane anesthesia. Monkeys were intubated, and a peripheral catheter was placed. Preemptive analgesia (buprenorphine 0.01 mg/kg, IM; meloxicam 0.2 mg/kg, SC) and antibiotics (penicillin-G or cefotaxime) were administered. A hemilamenictomy was performed to isolate the dura and a 3 french cathether was implanted subdurally in the lumbar spine at the L4:L5 junction. The cannula was connected to an MRI compatible subcutaneous port (SAI Infusion Technologies, Lake Villa, IL). The cannula was extended subcutaneousy and the port was placed in the proximal flank. Post-op care included monitoring and additional analgesia (buprenorphine and/or meloxicam every 8–12 h for 48+ h as needed).

### Preparation and administration of AβO

AβO were prepared and validated as previously described in Wakeman et al. 2022. Briefly, to ensure consistency, parallel oligomerization reactions were pooled, quantified, aliquoted, flash frozen, and stored at -80°C. AβO exhibited the characteristic mix of monomers, dimers and trimers after western blot similar to previous AβO synthesis (Wakeman et al., 2022; data not shown). There were 12 lumbar IT administrations over one month with 200 μg AβO per injection in young adult AGMs.

### CSF sample collection

CSF was sampled from anesthetized monkeys with the dorsal neck aseptically prepared and percutaneous needle inserted into the cisterna manga to collect approximately 1 mL of CSF. CSF was transferred to sterile tubes and centrifuged ay 1700 x g for ∼ 10 minutes (4°C) prior to freezing in the vapor phase of liquid nitrogen in aliquots of 200 µL.

### EVps Extraction

We extracted EVps from all six individuals. We used a modified version of the Wako MagCapture protocol to enable direct protein extraction to minimize protein loss associated with downstream desalting steps. All manufacturer recommended steps were followed up to the elution stage. Samples were received in 200 µL aliquots. To achieve a sufficient input volume, three 200 µL tubes and one 75 µL tube were pooled, yielding a total of 675 µL. The pooled sample was centrifuged at 10,000 rpm for 30 minutes at 4– 10 °C, and 625 µL of the supernatant was carefully transferred to avoid disturbing the pellet. EVps were then isolated using the MagCapture protocol. For elution, the standard buffer was replaced with a 50% trifluoroethanol (TFE) and 0.1 M ammonium bicarbonate solution to allow direct protein extraction compatible with LC-MS analysis. Elution was performed in two rounds using the same solution: 25 µL was added in the first round and incubated on a magnetic stand and the eluate collected. This was followed by a second elution with an additional 25 µL of the same solution. The eluate was pooled, and 12 µL of 5% acetic acid was added to acidify the combined sample. The final volume was divided into three 17.3 µL aliquots, with one aliquot used for downstream proteome analysis. Aliquots were flash-frozen in liquid nitrogen and lyophilized. After drying, the flask was backfilled with high-purity argon via HEPA-rated gas filters to establish an inert atmosphere devoid of proteinaceous contaminants.

### Mass Spectrometry

We performed proteomics on CSF (N=4) and EVps (N=6). Both whole CSF and EV samples were resuspended in a minimal volume of 1:1 tetrafluoroethylene and 0.1 M ammonium bicarbonate, reduced with 5 mM DTT at 25°C for 1 hr, and alkylated with 10 mM iodoacetamide at 25°C for 45 min protected from light. Samples were then diluted by 10x and digested with trypsin at 1:10 enzyme:protein (w/w, Promega #V5113) overnight at 37°C. No desalting was performed. Peptides were dried to completion, resuspended in Solvent A (0.1% formic acid in Fisher Optima LC/MS grade water), and quantified by absorbance at 280 nm. Injection amounts were normalized for all samples.

Samples were analyzed using a Thermo Scientific Ultimate 3000 RSLCnano ultra-high-performance liquid chromatography (UPLC) system coupled to a high-resolution Thermo Scientific Eclipse Tribrid Orbitrap mass spectrometer. Each sample was injected onto a nanoEase M/Z Peptide BEH C18 column (1.7 μm, 75 μm x 250 mm, Waters Corporation) and separated by reversed phase UPLC using a gradient of 4-30% Solvent B (0.1% formic acid in Fisher Optima LC/MS grade acetonitrile) over a 60-min gradient at 300 nL/min flow, followed by a ramp of 30-90% Solvent B over 30 min, for a 90-min total gradient. Peptides were eluted directly into the Eclipse using positive mode nanoflow electrospray ionization with a capillary voltage of 2200 V. MS1 scans were acquired at 120,000 resolution, with an AGC target of 4e5, maximum ion time set to Auto, RF lens setting of 30%, and a scan range of 300-1800 m/z. Data-dependent MS2 scans were acquired in the Orbitrap at a resolution of 15,000, for any ions over 5.0e4 intensity threshold, with AGC target set to Standard, maximum ion time set to Auto, mass range set to Normal, scan range set to Auto, isolation window set to 1.6 m/z, and a fixed cycle time of 3 sec. HCD fragmentation was applied with 30% energy. Dynamic exclusion was set to exclude for 30 sec after 1 observation, for charge states 2-8 only.

### Identification and quantification of proteins

To identify peptides and proteins from mass spectrometry-based proteomes data, we conducted a database search using the FragPipe platform with the MSFragger search engine^24^, using the proteome of the rhesus macaque (*Macaca mulatta*; Proteome ID: UP000006718) as the reference database. Additionally, to quantify protein abundance, we performed data dependent acquisition (DDA) label-free quantification using IonQuant^24^ within the FragPipe platform, applying default settings and normalization across runs.

Given that our samples were from green monkeys (*Chlorocebus sabaeus*), we also performed protein identification using the green monkey reference proteome and compared the identified proteins between the rhesus macaque and green monkey annotations (Figure S6). As the rhesus macaque proteome is more comprehensively annotated, we used protein identifications mapped to this reference proteome for GO analysis. Using the Rhesus macaque database yielded 278 identifed proteins, representing 83% of the proteins observed when using the AGM proteome.

## QUANTIFICATION AND STATISTICAL ANALYSIS

### Statistics and other analysis

To establish a reference for EV-derived proteins in our samples, we curated a list of proteins previously identified in CSF-derived EVps from prior studies (Chiasserini et al., 2014; Muraoka et al., 2020). This curated set was used to identify EV-enriched proteins present in both whole CSF and isolated EVps. We visualized unique and shared proteins among sample types using a Venn diagram.

Principal component analysis (PCA) was performed to assess whether samples clustered by sample type and treatment group, and to inform the selection of representative samples for downstream analyses. PCA was conducted using the prcomp function in R on scaled log₁₀-transformed protein intensity values. To ensure reliable input and avoid missing data across conditions, only proteins detected in at least 3 out of 4 CSF samples, and at least 4 out of 6 EVps samples were included.

To guide sample selection, PCA was performed separately for the CSF and EVps datasets. Based on their distance from the respective clusters, sample ABO_3 from the CSF group and samples ABO_3 and ABO_5 from the EVps group were excluded from further analysis (Figure S3). PCA was also conducted on the combined CSF and EVps dataset to visualize sample-level variation and potential group-specific clustering (Figure S3).

### Differential abundance analysis

We investigated treatment specific differences in proteome composition by analyzing CSF and EVps samples separately. Protein intensities were normalized across samples within Fragpipe. To ensure no condition was missing, we focused on proteins present in at least 3 out of 4 CSF samples or 4 out of 6 EVps samples. Global variation in protein composition was investigated using principal component analysis (PCA) on log₁₀-transformed intensity values.

To assess differential abundance and identify proteins significantly altered between AβO-treated and vehicle-treated groups in CSF or EVps samples, we first computed the log₁₀ fold change of mean values between the two groups (AβO-treated / vehicle-treated) for each protein, referred to as **log₁₀FC_mean**.

Given the limited sample size (4 CSF samples and 6 EVps samples), we also applied an edge effect analysis which captures the difference between the closest intensity values across treatment and control groups, defining **log₁₀FC_edge** as:

- **log₁₀FC_edge** = log10[(maximum of AβO-treated values) / (minimum of vehicle-treated values)], if log₁₀FC_mean < 0
- **log₁₀FC_edge** = log10 [(minimum of AβO-treated values) / (maximum of vehicle-treated values)], if log₁₀FC_mean > 0

Regulated Core group: If **log₁₀FC_mean** and **log₁₀FC_edge** share the same sign, this indicates that all AβO-treated samples consistently exhibit either higher or lower intensity values compared to vehicle-treated samples.

We defined upregulated proteins as those with log₁₀FC_mean > 0 and log₁₀FC_edge > 0, and downregulated proteins as those with log₁₀FC_mean < 0 and log₁₀FC_edge < 0.

### Gene Ontology (GO) enrichment analysis

For all proteins identified in CSF or EVps samples within treatment or control groups, we performed GO enrichment analysis of Cellular Component (CC) using an overrepresentation test implemented in the R package clusterProfiler (Yu et al., 2012). GO annotations for the Rhesus macaque organism were used. The top 30 enriched GO CC terms ranked by adjusted p-values, were visualized in a tree plot to show the hierarchical clustering of terms using the enrichplot package (Yu, 2025).

For differential abundant proteins, we performed GO enrichment analysis of Biological Process (BP) using the same approach. We ranked the proteins by log₁₀FC_mean. For CSF samples, we computed the enrichment for top 50% upregulated and top 50% downregulated proteins. For EVps samples, we choose the top 70% as the cutoff for upregulated and downregulated proteins in CSF and EVps samples separately. The top 20 GO terms ranked by adjusted p-values, were visualized in dot plots.

For upregulated and downregulated proteins in CSF and EVps samples separately, we performed Disease Ontology (DO) enrichment analysis using human homologous gene symbols in an overrepresentation test implemented in the R package DOSE (Yu et al., 2015). Consistent with GO analysis, we used the top 50% differential abundant proteins for CSF samples and the top 70% for EVp samples.

Additionally, we focused on proteins within the regulated core associated with Alzheimer’s disease (DOID:10652) and plotted their log₁₀ intensity values across samples to further assess differences in abundance between the two conditions.

## SUPPLEMENTAL INFORMATION

**Table S1. Protein-level quantification results generated using FragPipe.** The table reports protein identifiers (Protein.ID, Entry.Name, Gene), sequence attributes (Protein.Length, Organism, Protein.Existence, Description), probability metrics (Protein.Probability, Top.Peptide.Probability), and spectral evidence (peptide counts, total/unique/combined spectral counts). Spectral counts and intensity values are provided separately for each sample, along with MaxLFQ-normalized intensities. The column *Indistinguishable.Proteins* lists additional proteins not distinguished from the leading entry.

**Table 2. Enrichment statistics for Gene Ontology (GO) terms identified from proteins detected in FragPipe analysis.** Columns report the ratio of genes in the input set versus the background (GeneRatio, BgRatio), enrichment metrics (RichFactor, FoldEnrichment, zScore), and significance measures (p-value, adjusted p-value, q-value). The *geneID* column lists contributing genes, and *Count* indicates the number of genes associated with each term.

**Table S3. Protein-level log fold change estimates between AβO and vehicle conditions.** Columns include log fold mean and log10 fold changes from two approaches (*log10FC mean* and *log10FC edge*), alongside protein identifiers (Protein, Protein.ID, Entry.Name, Gene), sequence attributes (Protein Length, Organism, Protein Existence), and functional descriptions.

**Table S4. Enrichment results for functional categories for differentially abundant proteins.** Each row represents a significantly enriched pathway or ontology term (*Description*). Columns report the ratio of genes in the input set versus the background (GeneRatio, BgRatio), enrichment metrics (RichFactor, FoldEnrichment, zScore), and significance measures (p-value, adjusted p-value, q-value). The *geneID* column lists contributing genes, and *Count* indicates the number of genes associated with each term.

**Figure S5. Disease Ontology (DO) enrichment analysis for differentially abundant proteins.** Significantly enriched disease terms identified from proteins detected in FragPipe analysis. Each term (*Description*) is shown with enrichment metrics (GeneRatio, BgRatio, RichFactor, FoldEnrichment, zScore) and significance levels (p-value, adjusted p-value, q-value). The *geneID* column lists proteins associated with each disease term, and *Count* indicates the number of genes mapped.

---

**Figure S1.**
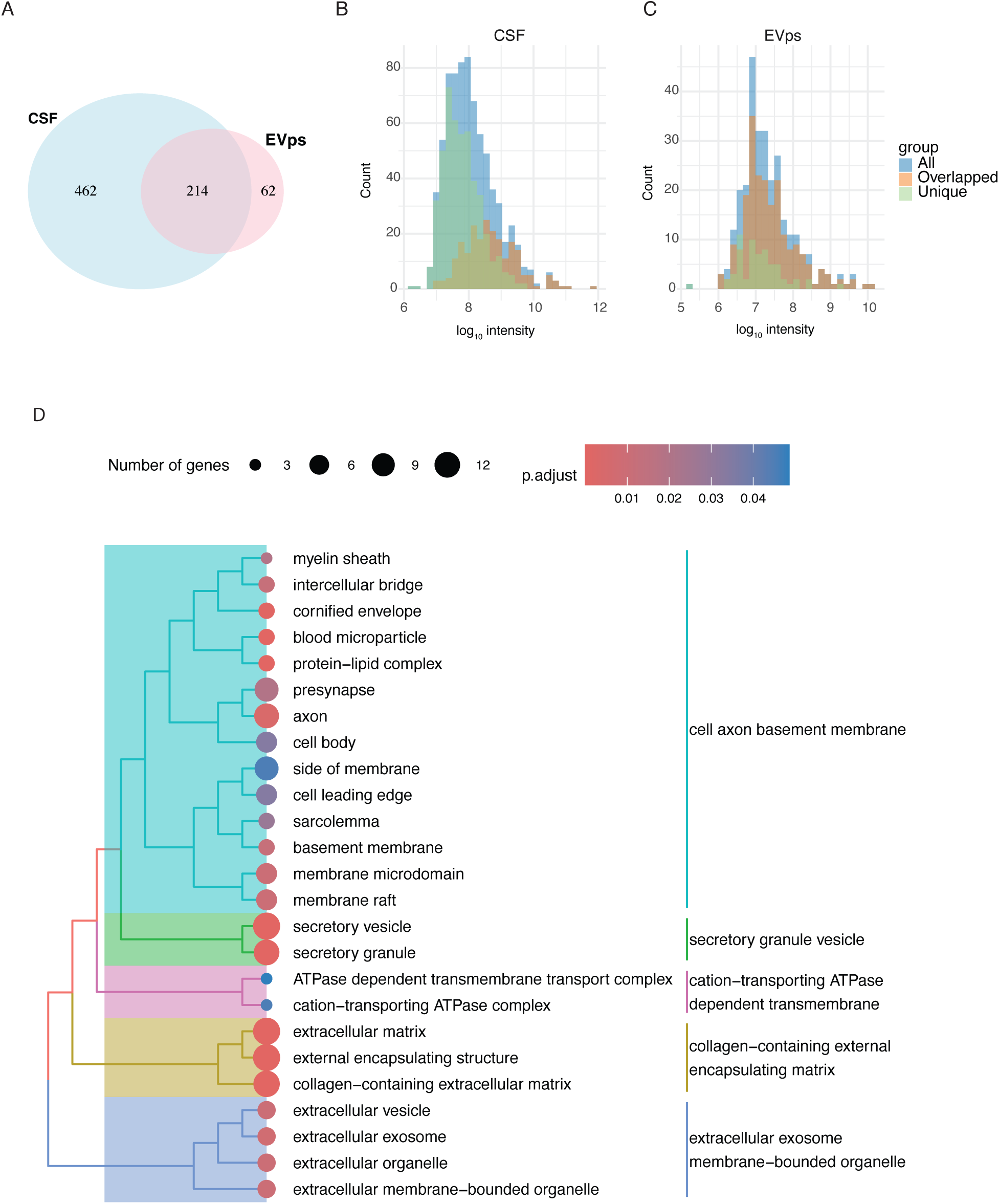
Overview of proteins across whole CSF and EVps. A. Overlap of proteins identified in CSF and EV samples. B-C. Comparison of the distribution of log₁₀-transformed median intensity for all proteins versus overlapping proteins and unique proteins (from A), shown separately for CSF (B) and EVps (C) samples. D. Top 30 enriched Gene Ontology (Cellular Component) terms ranked by adjusted p-values, and hierarchical clustering of similar terms, separately for CSF and EVps. See Figure S4 for a breakdown of protein identifications by sample and treatment category, and Figure S5 for Gene Ontology (GO) terms associated with biological processes and molecular functions.

**Figure S2.**
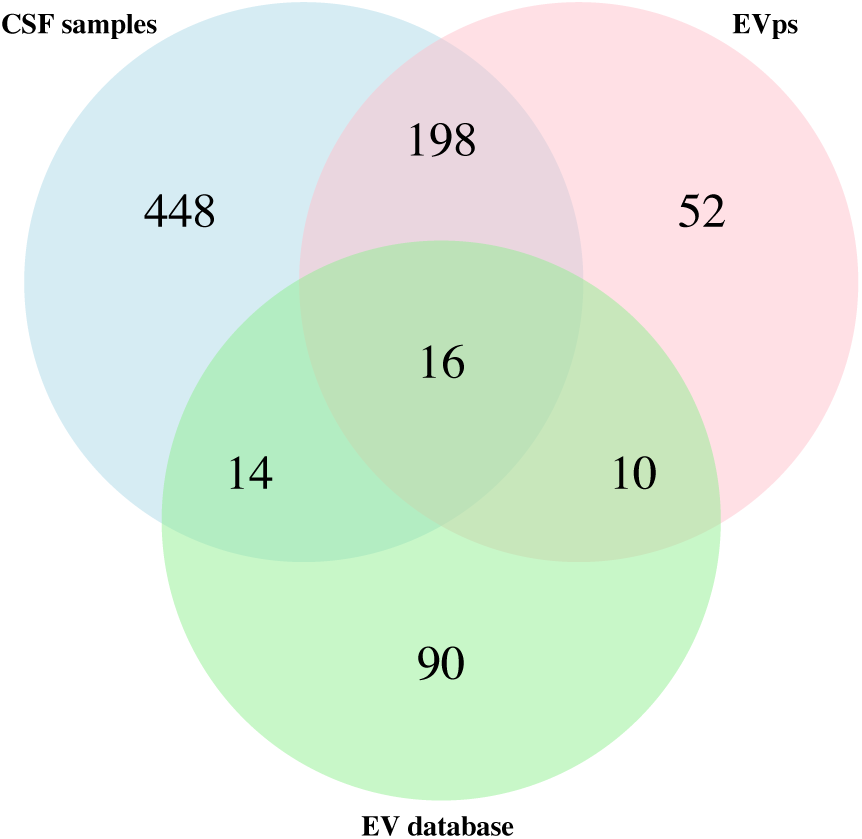
Comparison of our samples with published CSF EVps studies. The EVps reference databases are derived from human data, which may partially explain the low overlap. To verify the identity of our EVps preparations, we compared the EVps proteome to previously published EVps datasets. We found substantial overlap with canonical EVps proteins, supporting the effectiveness of our EVps enrichment strategy. Notably, the following proteins were detected in both the EVps and whole CSF fractions: APOA1, ANXA5, MBP, RAN, CFL1, NFASC, APOC3, HSPA8, DAG1, CLU, APOA2, HSPB1, APOD, PLTP, APOC2, and GFAP. While these proteins were present in both fractions, their relative abundances differed. Of the 16 overlapping proteins, 11 ranked within the top 50% in the EVps proteome, compared to 8 in the whole CSF proteome. We also identified 10 proteins that overlapped with known EVps markers but were uniquely detected in our EVps samples (ANXA4, ANXA7, ANXA1, ANXA11, CD9, ANXA2, CRYAB, ANXA6, PLP1, and NPTN). In contrast, 14 proteins were uniquely detected in the CSF proteome but also present in the EVps database, including VWF, AGRN, PFN2, ARF1, L1CAM, HSP90AA1, S100B, GLUL, APOB, APOA4, SNCB, SYN1, HSPH1, and MMP2.

**Figure S3.**
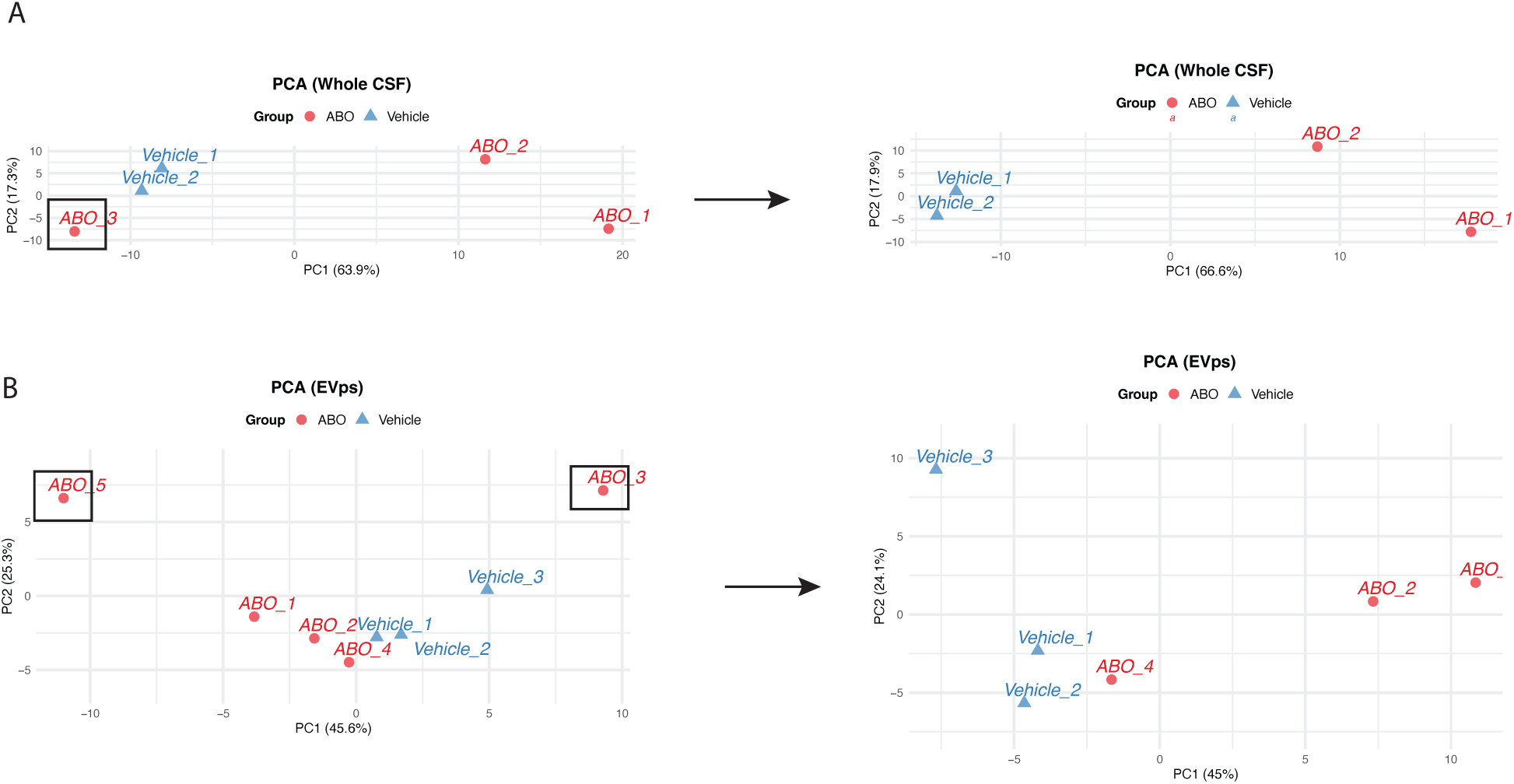
Global proteome differences between AβO-treated and vehicle-treated individuals. Principal component analysis (PCA) of log₁₀-transformed protein intensities from whole CSF (A) or EVps (B) samples, shown with all samples included (left) and with outliers removed (right).

**Figure S4.**
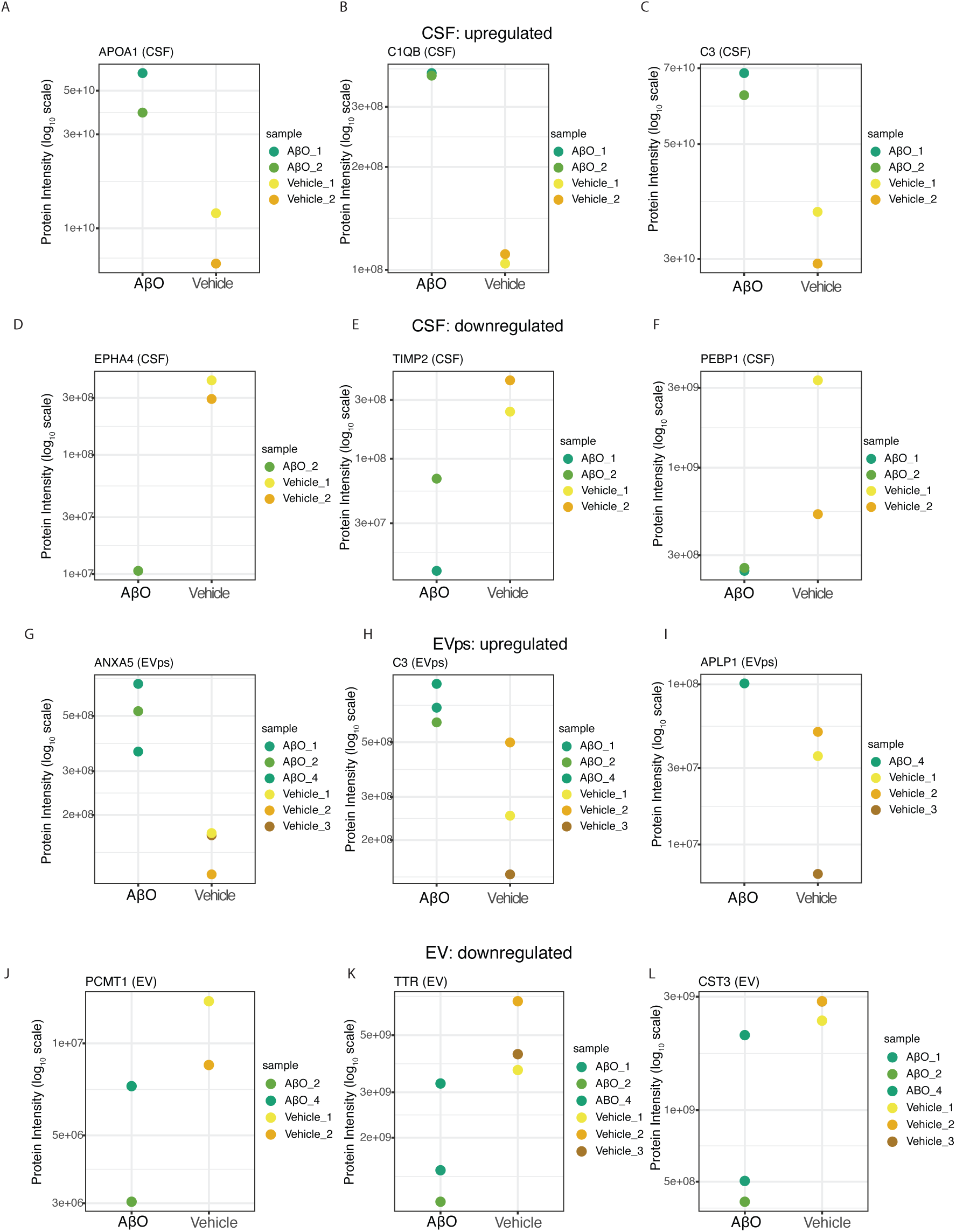
Selected AD-associated, differentially abundant proteins. Protein intensities of the most up- and down-regulated Alzheimer’s-associated proteins, shown with gene name for whole CSF (A–F) and EV compartments (G–L).

**Figure S5.**
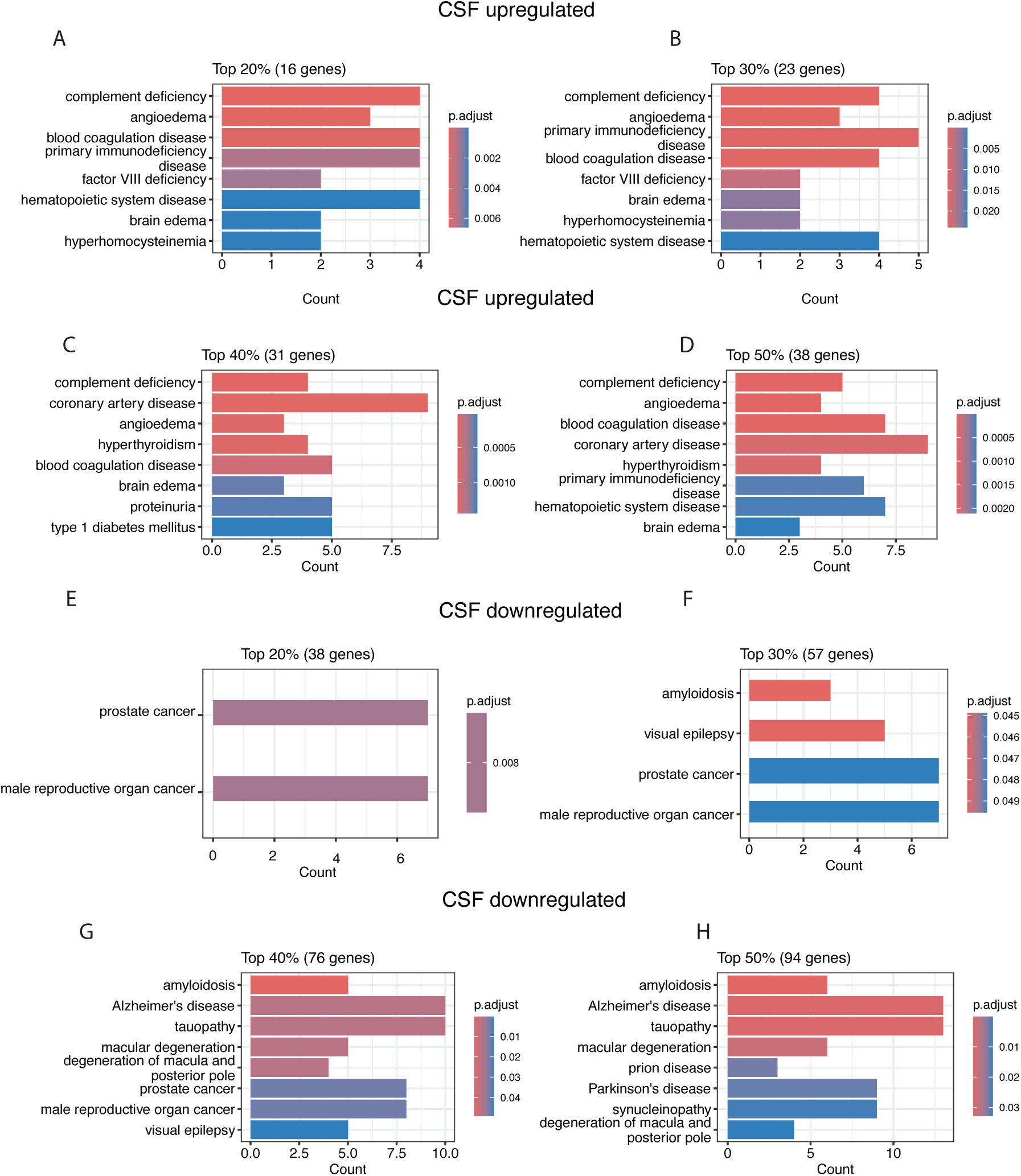
Disease Ontology (DO) Enrichment Analysis of Differentially Abundant Proteins in CSF. Bar plots display significantly enriched Disease Ontology (DO) terms (adjusted p-value < 0.05) for regulated core proteins in cerebrospinal fluid (CSF). Enrichment analyses were conducted separately for the top 20%, 30%, 40%, and 50% of (A-D) upregulated and (E-H) downregulated proteins. DO terms are ranked by adjusted p-value. The x-axis indicates the number of proteins associated with each enriched disease category.

**Figure S6.**
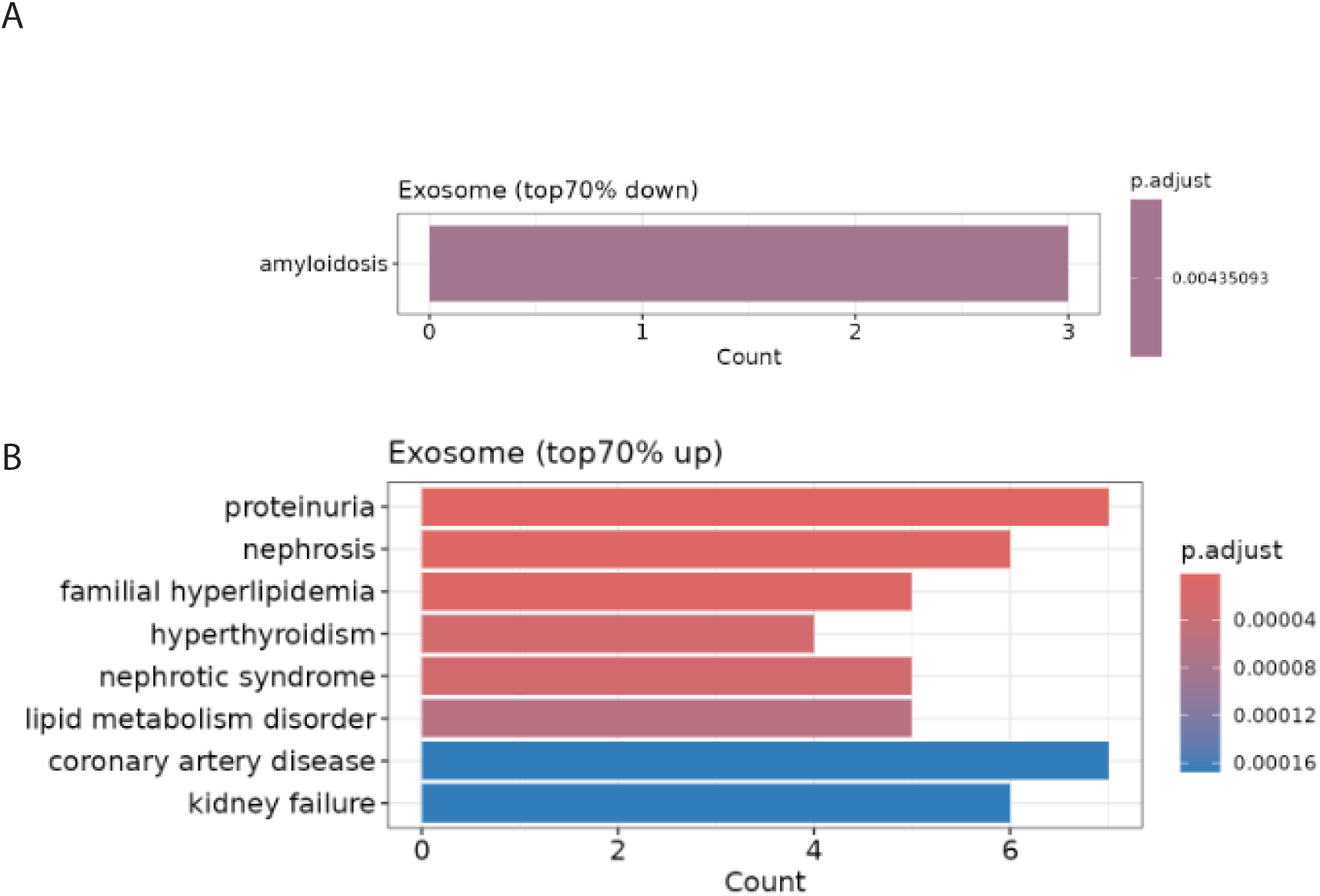
Disease Ontology (DO) Enrichment Analysis of Differentially Abundant Proteins in EVps. Bar plots display significantly enriched Disease Ontology (DO) terms (adjusted p-value < 0.05) for regulated core proteins in EVps. Enrichment analyses were conducted separately for the top 70% of (A) upregulated proteins and (B) downregulated. DO terms are ranked by adjusted p-value. The x-axis indicates the number of proteins associated with each enriched disease category.

## AUTHOR CONTRIBUTIONS

B.R.P.B. and A.D.M. conceptualized the study. M.R.W., J.D.E., and M.L. conducted the AGM experiment and sample collection. B.R.P.B., M.R.G., I.R.L., S.D., M.N.G., and A.D.M. performed the laboratory work. X.L. and G.G. led the analyses with input from B.R.P.B. and A.D.M. A.D.M. supervised the study. B.R.P.B. wrote the initial manuscript with input from X.L., A.D.M., and G.G. All authors contributed to the final version of the manuscript.

## ACKNOWLEDGEMENTS

This work was supported by the National Science Foundation Postdoctoral Fellowship awarded to BRPB, the Small Business Innovation Research grant AG067832 awarded to MW, the National Institute Office of Director grant R03OD036491 awarded to GG, and the National Institute on Aging grant R01AG068285 awarded to ADM. Special thanks to Jeremy Balsbaugh and Jen Liddle at the University Connecticut Proteomics & Metabolomics Facility, supported in part by NIH High-End Instrumentation Award 1S10-OD028445.

## Resource availability

### Lead contact

Further information and requests for resources should be directed to and will be fulfilled by the lead contact, Andrew Miranker.

### Data and code availability

- Data for this study are publicly available as of the date of publication. The mass spectrometry proteomics data have been deposited to the ProteomeXchange Consortium via the PRIDE partner repository with the dataset identifier PXD068600. Reviewer Token: ymHjjFlkv5CL
- All original code for data processing and performing downstream statistics can be found at https://github.com/G2Lab/NHP_Alzheimers_Proteomics
- Any additional information required to reanalyze the data reported in this paper is available from the lead contact upon request.

## REFERENCES

1. Hardy, J., and Selkoe, D.J. (2002). Medicine - The amyloid hypothesis of Alzheimer’s disease: Progress and problems on the road to therapeutics. Science 297, 353–356. DOI 10.1126/science.1072994.

2. Lanoiselée, H.M., Nicolas, G., Wallon, D., Rovelet-Lecrux, A., Lacour, M., Rousseau, S., Richard, A.C., Pasquier, F., Rollin-Sillaire, A., Martinaudl, O., et al. (2017). APP, PSEN1, and PSEN2 mutations in early-onset Alzheimer disease: A genetic screening study of familial and sporadic cases. Plos Med 14. e1002270 10.1371/journal.pmed.1002270.

3. Cline, E.N., Bicca, M.A., Viola, K.L., and Klein, W.L. (2018). The Amyloid-β Oligomer Hypothesis: Beginning of the Third Decade. Adv Alzh Dis 6, 565–608. 10.3233/978-1-61499-876-1-565.

4. De Strooper, B., and Karran, E. (2016). The Cellular Phase of Alzheimer’s Disease. Cell 164, 603–615. 10.1016/j.cell.2015.12.056.

5. Tung, J.Y., Lange, E.C., Alberts, S.C., and Archie, E.A. (2023). Social and early life determinants of survival from cradle to grave: A case study in wild baboons. Neurosci Biobehav R 152. 105282 10.1016/j.neubiorev.2023.105282.

6. Latimer, C.S., Shively, C.A., Keene, C.D., Jorgensen, M.J., Andrews, R.N., Register, T.C., Montine, T.J., Wilson, A.M., Neth, B.J., Mintz, A., et al. (2019). A nonhuman primate model of early Alzheimer’s disease pathologic change: Implications for disease pathogenesis. Alzheimers & Dementia 15, 93–105. 10.1016/j.jalz.2018.06.3057.

7. Forny-Germano, L., Silva, N.M.L.E., Batista, A.F., Brito-Moreira, J., Gralle, M., Boehnke, S.E., Coe, B.C., Lablans, A., Marques, S.A., Martinez, A.M.B., et al. (2020). Alzheimer’s Disease-Like Pathology Induced by Amyloid-β Oligomers in Nonhuman Primates (vol 34, pg 13629, 2014). J Neurosci 40, 8204–8204. 10.1523/Jneurosci.1827-20.2020.

8. Kumar, S., Stecher, G., Suleski, M., and Hedges, S.B. (2017). TimeTree: A Resource for Timelines, Timetrees, and Divergence Times. Molecular biology and evolution 34, 1812–1819. 10.1093/molbev/msx116.

9. Kalinin, S., Willard, S.L., Shively, C.A., Kaplan, J.R., Register, T.C., Jorgensen, M.J., Polak, P.E., Rubinstein, I., and Feinstein, D.L. (2013). Development of amyloid burden in African Green monkeys. Neurobiol Aging 34, 2361–2369. 10.1016/j.neurobiolaging.2013.03.023.

10. Jankowsky, J.L., and Zheng, H. (2017). Practical considerations for choosing a mouse model of Alzheimer’s disease. Mol Neurodegener 12. 89 10.1186/s13024-017-0231-7.

11. Drummond, E., and Wisniewski, T. (2017). Alzheimer’s disease: experimental models and reality. Acta Neuropathol 133, 155–175. 10.1007/s00401-016-1662-x.

12. Corey, T.M., Illanes, O., Lawrence, M., Perez, S.E., Liddie, S., and Callanan, J.J. (2023). Naturally occurring histological findings and Alzheimer’s-like pathology in the brain of aging African green monkeys (). J Comp Neurol 531, 1276–1298. 10.1002/cne.25494.

13. Wakeman, D.R., Weed, M.R., Perez, S.E., Cline, E.N., Viola, K.L., Wilcox, K.C., Moddrelle, D.S., Nisbett, E.Z., Kurian, A.M., Bell, A.F., et al. (2022). Intrathecal amyloid-beta oligomer administration increases tau phosphorylation in the medial temporal lobe in the African green monkey: A nonhuman primate model of Alzheimer’s disease. Neuropath Appl Neuro e1280010.1111/nan.12800.

14. Paolicelli, R.C., Bergamini, G., and Rajendran, L. (2019). Cell-to-cell Communication by Extracellular Vesicles: Focus on Microglia. Neuroscience 405, 148–157. 10.1016/j.neuroscience.2018.04.003.

15. Chiasserini, D., van Weering, J.R.T., Piersma, S.R., Pham, T.V., Malelnadeh, A., Teunissen, C.E., de Wit, H., and Jiménez, C.R. (2014). Proteomic analysis of cerebrospinal fluid extracellular vesicles: A comprehensive dataset. J Proteomics 106, 191–204. 10.1016/j.jprot.2014.04.028.

16. Del Campo, M., Peeters, C.F.W., Johnson, E.C.B., Vermunt, L., Hok, A.H.Y.S., van Nee, M., Chen-Plotkin, A., Irwin, D.J., Hu, W.T., Lah, J.J., et al. (2022). CSF proteome profiling across the Alzheimer’s disease spectrum reflects the multifactorial nature of the disease and identifies specific biomarker panels. Nat Aging 2, 1040–1053. 10.1038/s43587-022-00300-1.

17. Visser, P.J., Reus, L.M., Gobom, J., Jansen, I., Dicks, E., van der Lee, S.J., Tsolaki, M., Verhey, F.R.J., Popp, J., Martinez-Lage, P., et al. (2022). Cerebrospinal fluid tau levels are associated with abnormal neuronal plasticity markers in Alzheimer’s disease (vol 17, pg 1, 2022). Mol Neurodegener 17. 37 10.1186/s13024-022-00540-0.

18. Higginbotham, L., Ping, L.Y., Dammer, E.B., Duong, D.M., Zhou, M.T., Gearing, M., Hurst, C., Glass, J.D., Factor, S.A., Johnson, E.C.B., et al. (2020). Integrated proteomics reveals brain-based cerebrospinal fluid biomarkers in asymptomatic and symptomatic Alzheimer’s disease. Sci Adv 6. eaaz9360 10.1126/sciadv.aaz9360.

19. Ali, M., Western, D., Liu, M.H., Beric, A., Budde, J., Do, A., Heo, G., Wang, L.H., Gentsch, J., Schindler, S.E., et al. (2025). Multi-cohort cerebrospinal fluid proteomics identifies robust molecular signatures across the Alzheimer disease continuum. Neuron 113. 10.1016/j.neuron.2025.02.014.

20. Bader, J.M., Geyer, P.E., Müller, J.B., Strauss, M.T., Koch, M., Leypoldt, F., Koertvelyessy, P., Bittner, D., Schipke, C.G., Incesoy, E.I., et al. (2020). Proteome profiling in cerebrospinal fluid reveals novel biomarkers of Alzheimer’s disease. Mol Syst Biol 16. e9356 10.15252/msb.20199356.

21. Muraoka, S., Jedrychowski, M.P., Yanamandra, K., Ikezu, S., Gygi, S.P., and Ikezu, T. (2020). Proteomic Profiling of Extracellular Vesicles Derived from Cerebrospinal Fluid of Alzheimer’s Disease Patients: A Pilot Study. Cells-Basel 9. 1959 10.3390/cells9091959.

22. Chatterjee, M., Oezdemir, S., Kunadt, M., Koel-Simmelink, M., Boiten, W., Piepkorn, L., Pham, T.V., Chiasserini, D., Piersma, S.R., Knol, J.C., et al. (2023). C1q is increased in cerebrospinal fluid-derived extracellular vesicles in Alzheimer’s disease: A multi-cohort proteomics and immuno-assay validation study. Alzheimers Dement 19, 4828–4840. 10.1002/alz.13066.

23. Hirschberg, Y., Valle-Tamayo, N., Dols-Icardo, O., Engelborghs, S., Buelens, B., Vandenbroucke, R.E., Vermeiren, Y., Boonen, K., and Mertens, I. (2023). Proteomic comparison between non-purified cerebrospinal fluid and cerebrospinal fluid-derived extracellular vesicles from patients with Alzheimer’s, Parkinson’s and Lewy body dementia. J Extracell Vesicles 12. e12383 10.1002/jev2.12383.

24. Yu, F.C., Teo, G.C., Kong, A.T., Fröhlich, K., Li, G.X., Demichev, V., and Nesvizhskii, A.I. (2023). Analysis of DIA proteomics data using MSFragger-DIA and FragPipe computational platform. Nat Commun 14. 10.1038/s41467-023-39869-5.

25. Blennow, K., Hampel, H., Weiner, M., and Zetterberg, H. (2010). Cerebrospinal fluid and plasma biomarkers in Alzheimer disease. Nat Rev Neurol 6, 131–144. 10.1038/nrneurol.2010.4.

26. Santandrea, E., Aliakbari, F., Truscott, E., Mccaig, L., Donison, N.S., Graham, D., Strong, M.J., and Volkening, K. (2024). A technique for repeated blood and cerebrospinal fluid sampling from individual rats over time without the need for repeated anesthesia. Sci Rep-Uk 14. 517110.1038/s41598-024-55666-6.

27. Oshio, K., Watanabe, H., Song, Y., Verkman, A.S., and Manley, G.T. (2004). Reduced cerebrospinal fluid production and intracranial pressure in mice lacking choroid plexus water channel Aquaporin-1. Faseb J 18, 76–+. 10.1096/fj.04-1711fje.

28. Association of Primate Veterinarians Guidelines for Cerebrospinal Fluid Aspiration for Nonhuman Primates in Biomedical Research. (2024). J Am Assoc Lab Anim 63, 357–358.

29. Tomiyama, T., Matsuyama, S., Iso, H., Umeda, T., Takuma, H., Ohnishi, K., Ishibashi, K., Teraoka, R., Sakama, N., Yamashita, T., et al. (2010). A Mouse Model of Amyloid β Oligomers: Their Contribution to Synaptic Alteration, Abnormal Tau Phosphorylation, Glial Activation, and Neuronal Loss. J Neurosci 30, 4845–4856. 10.1523/Jneurosci.5825-09.2010.

30. Baerends, E., Soud, K., Folke, J., Pedersen, A.K., Henmar, S., Konrad, L., Lycas, M.D., Mori, Y., Pakkenberg, B., Woldbye, D.P.D., et al. (2022). Modeling the early stages of Alzheimer’s disease by administering intracerebroventricular injections of human native Aβ oligomers to rats. Acta Neuropathologica Communications 10. 113 10.1186/s40478-022-01417-5.

31. Gomes, P., Tzouanou, F., Skolariki, K., Vamvaka-Iakovou, A., Noguera-Ortiz, C., Tsirtsaki, K., Waites, C.L., Vlamos, P., Sousa, N., Costa-Silva, B., et al. (2022). Extracellular vesicles and Alzheimer’s disease in the novel era of Precision Medicine: implications for disease progression, diagnosis and treatment. Exp Neurol 358. 114183 10.1016/j.expneurol.2022.114183.

32. Iturria-Medina, Y., Sotero, R.C., Toussaint, P.J., Mateos-Pérez, J.M., Evans, A.C., and Neuroimaging, A.s.D. (2016). Early role of vascular dysregulation on late-onset Alzheimer’s disease based on multifactorial data-driven analysis. Nat Commun 7. 1193410.1038/ncomms11934.

33. Heneka, M.T., Carson, M.J., El Khoury, J., Landreth, G.E., Brosseron, F., Feinstein, D.L., Jacobs, A.H., Wyss-Coray, T., Vitorica, J., Ransohoff, R.M., et al. (2015). Neuroinflammation in Alzheimer’s disease. Lancet Neurol 14, 388–405. Doi 10.1016/S1474-4422(15)70016-5.

34. Cirrito, J.R., Yamada, K.A., Finn, M.B., Sloviter, R.S., Bales, K.R., May, P.C., Schoepp, D.D., Paul, S.M., Mennerick, S., and Holtzman, D.M. (2005). Synaptic activity regulates interstitial fluid amyloid-β levels in vivo. Neuron 48, 913–922. 10.1016/j.neuron.2005.10.028.

35. Végh, M.J., Heldring, C.M., Kamphuis, W., Hijazi, S., Timmerman, A.J., Li, K.W., van Nierop, P., Mansvelder, H.D., Hol, E.M., Smit, A.B., and van Kesteren, R.E. (2014). Reducing hippocampal extracellular matrix reverses early memory deficits in a mouse model of Alzheimer’s disease. Acta Neuropathol Com 2. 76 10.1186/s40478-014-0076-z.

36. Panyard, D.J., McKetney, J., Deming, Y.K., Morrow, A.R., Ennis, G.E., Jonaitis, E.M., Van Hulle, C.A., Yang, C.R., Sung, Y.J., Ali, M., et al. (2023). Large-scale proteome and metabolome analysis of CSF implicates altered glucose and carbon metabolism and succinylcarnitine in Alzheimer’s disease. Alzheimers Dement 19, 5447–5470. 10.1002/alz.13130.

37. Stoye, N.M., Jung, P., Guilherme, M.D., Lotz, J., Fellgiebel, A., and Endres, K. (2020). Apolipoprotein A1 in Cerebrospinal Fluid Is Insufficient to Distinguish Alzheimer’s Disease from Other Dementias in a Naturalistic, Clinical Setting. J Alzheimers Dis Rep 4, 15–19. 10.3233/Adr-190165.

38. Xie, G.F., Jiang, G.G., Huang, L.Q., Sun, S.Q., Wan, Y.W., Li, F., Wu, B.J., Zhang, Y., Li, X.Y., Xiong, B.W., and Xiong, J. (2025). The Role of APOA-I in Alzheimer’s Disease: Bridging Peripheral Tissues and the Central Nervous System. Pharmaceuticals-Base 18. 790 10.3390/ph18060790.

39. Hirschberg, Y., Boonen, K., Schildermans, K., van Dam, A., Pintelon, I., Vandendriessche, C., Velimirovic, M., Jacobs, A., Vandenbroucke, R.E., Nelissen, I., et al. (2022). Characterising extracellular vesicles from individual low volume cerebrospinal fluid samples, isolated by SmartSEC. J Extracell Biol 1. e55 10.1002/jex2.55.

